# Non-destructive assessment of multi-material micro-tissue mechanics reveals the critical role of rigidity gradients in tumour growth and pressure

**DOI:** 10.1101/2025.05.14.653936

**Authors:** Angélique Ala, Camille Douillet, Audrey Ferrand, Vincent Velay, Stéphane Segonds, Gaëlle Recher, Florian Bugarin

## Abstract

Probing stiffness anisotropies in three-dimensional materials non-destructively is a major challenge in disciplines as diverse as aeronautics and medicine. While the former typically relies on various mechanical tests—such as tensile, compression, bending, and shear—performed on sample parts, the latter often employs acoustic techniques or wave propagation through matter. The choice of techniques will depends on the size of the sample of interest and the desired resolution. In our case, to probe the mechanical properties of sub-millimetre micro-tissues, it is necessary to use methods with high resolution and as furtive as possible. We present a method based, 1/ on imaging the displacement of microbeads within a hydrogel resulting from the growth of a three-dimensional micro-tissue and, 2/ on finite element modelling of the deformations underlying bead displacements. This approach allows us to determine the elastic properties of the hydrogel and, in particular, to show that beyond a certain thickness, incomplete cross-linking of the hydrogel results in a stiffness gradient. We show that when the micro-tissue contacts with an immediately rigid alginate wall, the pressure exerted over time increases very rapidly, whereas when the micro-tissue encounters a substrate with a stiffness gradient, the pressure increase is more gradual. Uncovering this could provide a better understanding of the role of tumour microenvironment stiffness in metastatic escape processes.

## 1. Introduction

Whether natural (embryos, organs or tumour) or synthetic, i.e. bio-engineered (spheroids, organoids, bio-printed tissues), 3D multicellular assemblies are embedded in an enveloping environment. This environment may be neighbouring cells, soft or stiff tissues or hydrogels when cultured *in vitro* (Matrigel, alginate, GelMA, collagen[1,2]). Mechanical interactions within the tissue itself influence collective cell behaviour (“go or grow” behaviour)[2,3] or intercellular signalling pathways[4]. Furthermore, the tissue also interacts with its environment, applying stresses to it. Due to the principle of reciprocity of mechanical actions, both the tissue of interest and its surrounding environment experiences the same stress. Therefore cells adapt permanently their behaviour to any change in their immediate environment[4–7]. Quantifying these mechanical interactions would help to decipher the interplay between the tissue of interest and its surroundings, whether under physiological or pathological conditions. This would allow us to understand how cells modify their behaviour under pressure and what are the consequences of losing this reciprocity of mechanical actions[8,9]. Several methods are available to map the forces and stresses between the tissue of interest and its environment. First, those applicable to 2D cultures include 2D Traction Force Microscopy (TFM)[10–12], micropillar arrays[13,14], or monolayer stress microscopy[15,16]. It is also possible to map mechanical stresses in volume, either using 3D TFM[17–19], which provides 3D traction forces from the displacement of the Extra-Cellular Matrix (ECM); or the micro bulge test[20], which relies on the formation of domes whose deformations are measured. Another approach involves inserting deformable but incompressible oil droplets within tissues, like zebrafish embryos, and monitoring their deformation to extract forces applied[21]. However, these methods and their implementation have limitations. They are less applicable when the ECM is not purely elastic. Thus, it can be extremely challenging to recover forces associated with ECM deformations. Ideally, to extract quantitative information with biological relevance, it is necessary to provide a tuneable and reproductible system that allows all-optical measurement of a high number of specimens. To this aim, we use the Cellular Capsule Technology (CCT) that consists in encapsulating a cell suspension in hollow alginate spherical capsules[22–24] or tubes[25,26]. If fluorescent microbeads are incorporated into the non-reticulated alginate, upon jellification, the beads are trapped within the hydrogel mesh. The cells later grow, leading the cellular aggregate to eventually deform the alginate shell, leading the beads displacing according to the gel’s strain. By monitoring this process with 3D+time fluorescence confocal microscopy, it is possible to track the displacements of the beads. To retrieve quantitative information from these displacements, some hypotheses must be made regarding the mechanical properties of the material of interest (elasticity, viscoelasticity or plasticity), and the capsule’s shape and size. Alginate hydrogel has been extensively studied through rheology approaches[27–30], elongation measurements[31], or osmotic swelling measurements[22]. Depending on the deformation regime, alginate expresses non-linear responses[22,27], requiring a method that accounts for its material properties. Finite Element Modelling (FEM)[32] is particularly well-suited approach to this aim. This modelling method is used to represent and model the deformations and stresses of complex structures when there is no satisfying analytical solution[33–35]. FEM can model volumetric environments and provide both global and local information in 3D[36]. In this work, we present how we use of CCT to produce a high number of encapsulated fluorescent cellular aggregates, proceed to 3D+time live imaging of the micro-tissue growth over time and resulting capsule deformation. These experimental datasets are then used to design a custom-made FEM framework to retrieve strains and pressures applied by the micro-tissue to the capsule. We consider different geometrical features of the capsules and found out that not only we can predict pressure, but also deduce, model and test new hypothesis to the alginate material itself in the regime of large deformations. Ultimately, we found out that the pressure applied and sensed by the micro-tissue to its immediate environment directly depends on the stiffness of the material, the stiffer, the higher pressure rises (steep slope 3.108kPa). Whereas, when the alginate shell is thicker and exhibit a stiffness gradient with low stiffness in contact with the cells, the resulting pressure is lower (steep slope 0.867kPa). Here we provide an optical and digital pipeline to quantify mechanical properties of soft materials together with quantification of resulting pressures between the two components of the micro-tissue.

## 2. Results

### 2.1. Experimental based Finite Element Model and implementation of an inverse method to monitor volumetric deformation and compute tissue exerted pressure

The CCT[22,23] is a microfluidic technique developed to encapsulate cells and ECM inside a hollow alginate shell. The device consists of a 3D-printed microfluidic chip from which solutions are extruded concentrically[23] (Figure 1A). The outermost solution is a mix of 2% alginate with dispersed 1µm diameter fluorescent microbeads (1%, corresponding to 4.8 x10^5^ beads/mL, Sicastar, RedF, see Material & Methods section). The intermediate sorbitol solution (200mM) helps preventing alginate reticulation inside the chip. The innermost solution is the cell suspension, made of a mix of fluorescently labelled (LifeAct::eGFP) Caco2 cells and ECM (100 000 cells in 200µL DMEM and 200µL CultrexBME, see Material & Methods section). The jet breaks in the air and the capsules land in a calcium bath (100mM), where gelling occurs almost instantaneously. Alginate is a hydrogel that is porous to nutrients and optically transparent, making the capsules amenable for light microscopy. Deformations induced by cells to the alginate shell can be monitored and quantified, providing insights into the pressure exerted by the encapsulated tissue. Bright field time-lapse microscopy (Figure 1B) permits imaging and measuring shell deformation in the focal plane using detection algorithm[22]. However, these analyses are limited to the equatorial plane and roughly describe the average deformation, thus, missing possible spatial anisotropy mapping. The addition of fluorescent beads in the alginate, combined with 3D time-lapse microscopy provides 4D datasets of both the cell aggregate growth and the resulting deformation of the alginate shell (Figure 1C). The aim of this section is to describe our approach to calibrate the parameters of a Finite Element (FE) model based on the behaviour of the capsule subjected to the growth of cellular aggregates. This approach involves implementing three key steps: (i) acquiring experimental data, (ii) creating a geometric model to simulate the observed real system, and (iii) updating the model parameters through optimization by comparing experimental and numerical data.

**Figure 1.**
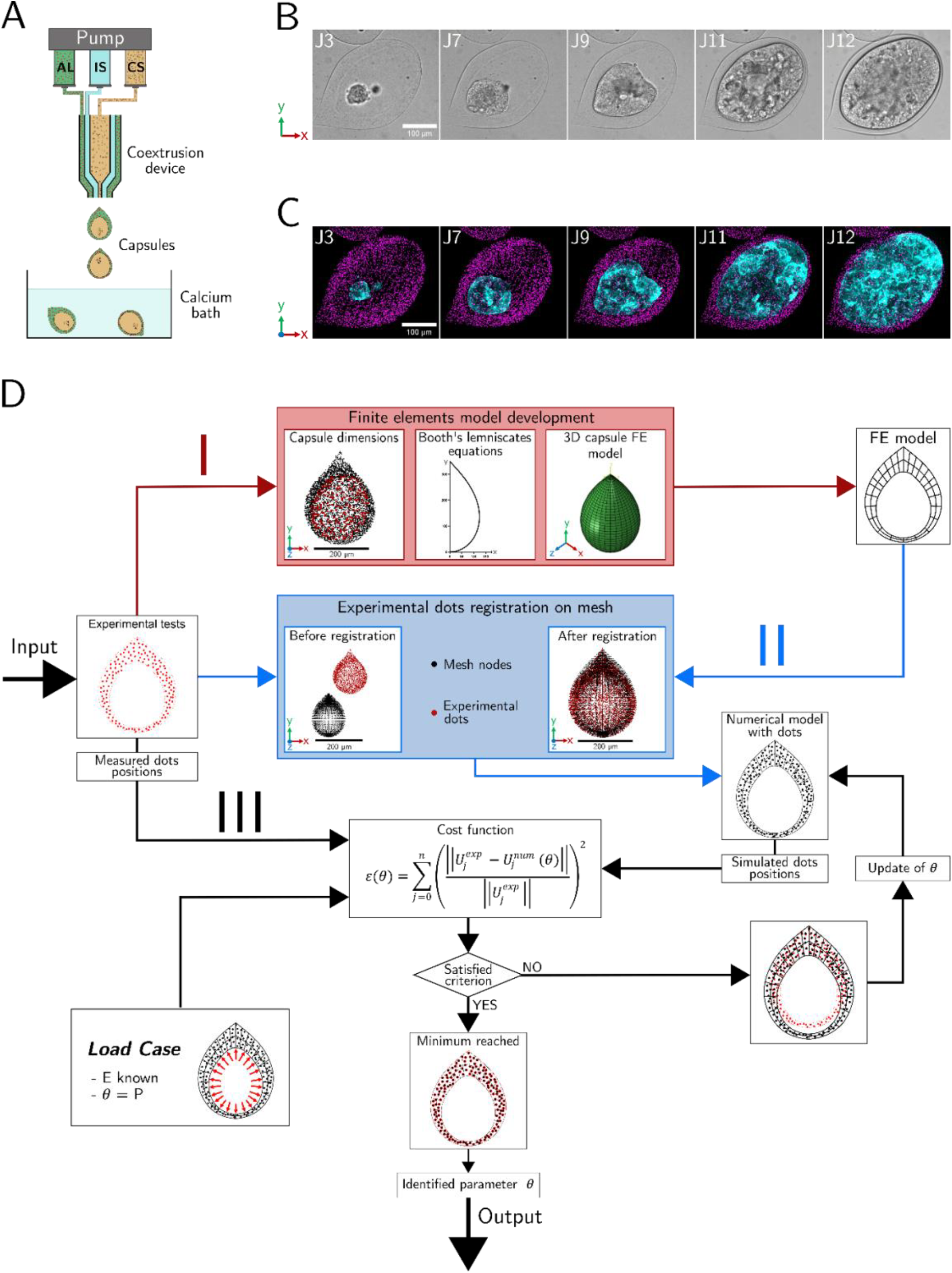
Caco2 filled alginate capsules 3D+time imaging and diagram of the inverse method used to compute internal pressure. (A) Schematic of the CCT. View of the microfluidic chip with the three solutions: alginate (AL) and fluorescent beads; intermediate sorbitol solution (IS) and the cell suspension (CS). After extrusion, capsules gel in the calcium bath. (B) Bright Field snapshots of encapsulated Caco2 cells over 12 days. (C) Corresponding fluorescent confocal maximum intensity projections of 3D z-stacks (bead in magenta and tissue labelled with LifeAct::eGFP actin in cyan). (D) Schematic of the inverse method used. The FE model is built by taking dimensions on the experimental point cloud (dots position) and by using the Booth lemniscate’s equations. Experimental data are registered on the mesh with an ICP registration. The aim of the inverse method is to find *θ* by minimizing *ε* with Nelder Mead algorithm.

The first step (i) involves tracking the displacements of fluorescent beads embedded in the alginate capsule using a 3D microscopy system and ImageJ software (Fijiyama[37] and Trackmate[38] plugins). This allows for the mapping of the bead displacement field resulting from the pressure exerted by the growing cellular aggregates on the capsule.

To achieve step (ii), we developed an FE model representing the capsule geometry with an outer radius ranging from 100 µm to 300 µm[24]. The initial dimensions (outer/inner heights, outer/inner widths, and capsule bottom thickness) of this model are measured from the initial positions (i.e., before deformation) of the fluorescent beads (Figure 1D, part I – step 1). These measurements then determine the mathematical parameters of a Booth lemniscate (see definition in Materials & Methods section: Virtual Model Geometry) representing the 2D profile of the capsule (Figure 1D, part I – step 2). The final FE model is obtained by revolving this 2D profile around the Y-axis and creating the finite elements using a Python algorithm (Figure 1D, part I – step 3). By imposing a displacement on the nodes of the FE mesh located on the inner wall of the capsule, we can simulate the pressure exerted by the cellular aggregate inside the capsule over time.

The third step (iii) (Figure 1D part III) can be divided into two phases: writing an error function, representing the discrepancy between the numerical model and the observations, and the process of minimizing this error by updating the model parameters.

The writing of the error function, denoted as ε, is based on the normalized comparison of the n positions of the n microbeads. Let 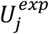 be the experimental position of bead j and 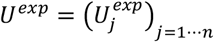 be the set of experimental positions of all n beads. Similarly, let 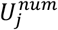 be the numerically simulated position of bead j and 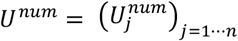 be the set of numerically simulated positions of all n beads. The first phase involves aligning the observations and the FE model of the capsule in the same 3D reference frame using an ICP algorithm[39] between the mesh nodes and the 3D positions of the beads (Figure 1D part II). By associating each bead with the centroid of a finite element, we obtain a set of experimental *U*^*exp*^ and simulated *U*^*num*^ 3D points in the same 3D reference frame. The second phase involves introducing a dependency parameter into each cloud *U*^*num*^ to update the numerical model. As a first approach, we can consider the triplet of parameters E (Young’s modulus), ν (Poisson’s ratio), and P (internal pressure) as the optimization variable θ. Indeed, numerically, the simulated positions of the microbeads *U*^*num*^ directly depend on the pressure P applied to the internal wall by the confluent tissue, as well as the Poisson’s ratio ν and the Young’s modulus of the alginate E. However, optimizing with the parameter θ containing three quantities (E, ν, and P) is very challenging due to the complexity of the models used. We assume that the pressure begins to exert itself as soon as the tissue reaches confluence, i.e., when it is in complete contact with the internal surface of the capsule. The Poisson’s ratio can be set to 0.5, as alginate is considered an incompressible material. We then conducted an experimental plan to evaluate the influence of E and P on the variability of the maximum Von Mises stress[40] (quantifying the intensity of stresses experienced by the capsule) on *U*^*num*^ to verify if it is possible to further reduce the size of the optimization parameter θ. Simulations were performed with values varying by ±30% around *E* = 94*kPa* and *P* = 4 *kPa*, which are the extreme values presented in Table 2B. We then carried out four simulations to calculate the numerical positions of the beads *U*^*num*^, combining the minimum and maximum values of E and P, and then calculated 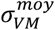 the average (in kPa) of the maximum Von Mises stress. The results (cf. Figure 2C) indicate that an increase in Young’s modulus from 66 *kPa* to 122 *kPa* leads to a 13% decrease in 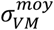, while a variation in pressure from 2.81 *kPa* to 5.19 *kPa* results in a 185% increase. The impact of P on the variability of *U*^*num*^ is thus approximately 12 times greater than that of E. It appears, therefore, that it is possible to set the Young’s modulus E to a given estimated value without introducing bias into the simulation results, as its influence is negligible compared to that of the pressure. Through osmotic pressure swelling measurements (see Material & Methods section), the Young’s modulus is set to *E* = 94 *kPa* in this model. The optimization parameter θ is thus limited to the pressure P, which constitutes our sole decision variable, as its variation leads to different deformed meshes and, consequently, different point clouds *U*^*num*^(*P*). The error function *ε*(*P*) is then defined as the root mean square, over the n time steps j, of the normalized discrepancies between the experimental *U*^*exp*^ and simulated *U*^*num*^(*P*) positions:

**Figure 2.**
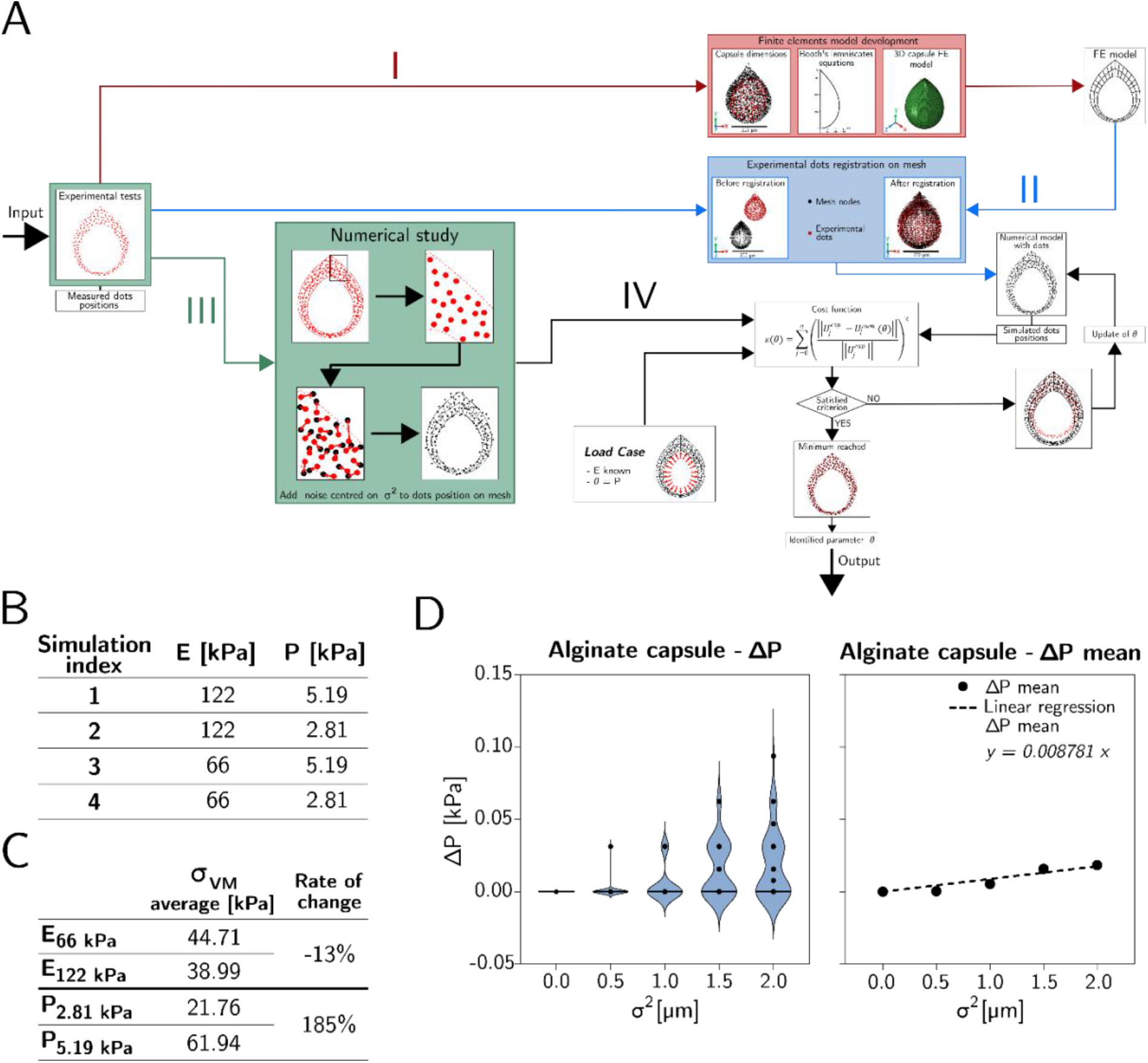
Computing numerical analysis reveals the importance of pressure over Young’s modulus. (A) Schematic of the inverse method with the numerical analysis step. (B) Values of input parameters used for the design experiment. (C) Variation of Von Mises (VM) stress according to Young’s modulus and pressure values for alginate capsule model. (D) Study of the noise sensibility of the alginate capsule model when pressure varies. Left: violin plot of the distribution of results. On the right: averaged results and the slope of the linear regression, which characterize the cost function sensitivity to noise.

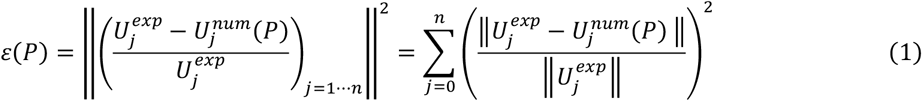

A minimization algorithm applied to *ε*(*P*) finally allows obtaining the value of the pressure 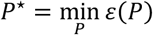 for which the numerical model best simulates the actual displacements of the microbeads. The algorithm used is the Nelder-Mead algorithm[41] with *P*_0_ = 0 as the initial guess

### 2.2. FEM Capsule Model Validation with Numerical Analysis

To numerically evaluate the robustness of the inverse method, we conducted an analysis to estimate the sensitivity of our approach to measurement noise arising from the tracking of experimental microbeads. For this purpose, we used a reference pressure of *P*^*ref*^ = 5*kPa* (with the Young’s modulus still fixed at *E* = 94 *kPa*) and generated, from the initial observed positions of the microbeads, a synthetic cloud of n positions 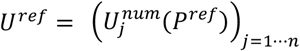. Noise was then added to the coordinates of the cloud *U*^*ref*^ according to a Gaussian distribution centered at 0 with different variance values *σ*^2^ ([0.0, 0.5, 1.0, 1.5, 2.0] in *μm*). This resulted in a cloud 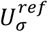 which we substituted for our experimental cloud *U*^*exp*^ in (1) to obtain a discrepancy function *ε*_*σ*_(*P*). Minimizing this function allowed us to obtain a pressure 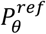 and the associated error 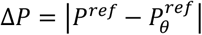, quantifying the influence of noise on our inverse method. For each variance value, we generated 100 simulations of the cloud 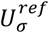, calculated the corresponding pressure values 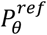, averaged them, and plotted them as a function of *σ*^2^ in Figure 2D. The results show that the slope of the trend line is less than 0.01, which is 0.2% of *P*_*ref*_, indicating very low sensitivity to noise. This demonstrates that the proposed inverse method is particularly numerically robust to measurement noise related to the extraction of the 3D positions of the microbeads.

### 2.3. Pressure exerted on the inner wall of alginate shells by cell aggregates

With our model numerically validated, we conducted a study on six different capsules, identified as A to F (see Figure 3). To standardize the processing scales, we define, for each alginate capsule, a reference time *T*_0_ corresponding to the last acquisition moment before confluence, at which the internal pressure is assumed to be zero. The uniaxial material law for alginate, assumed to be isotropic and homogeneous, is as follows:

**Figure 3.**
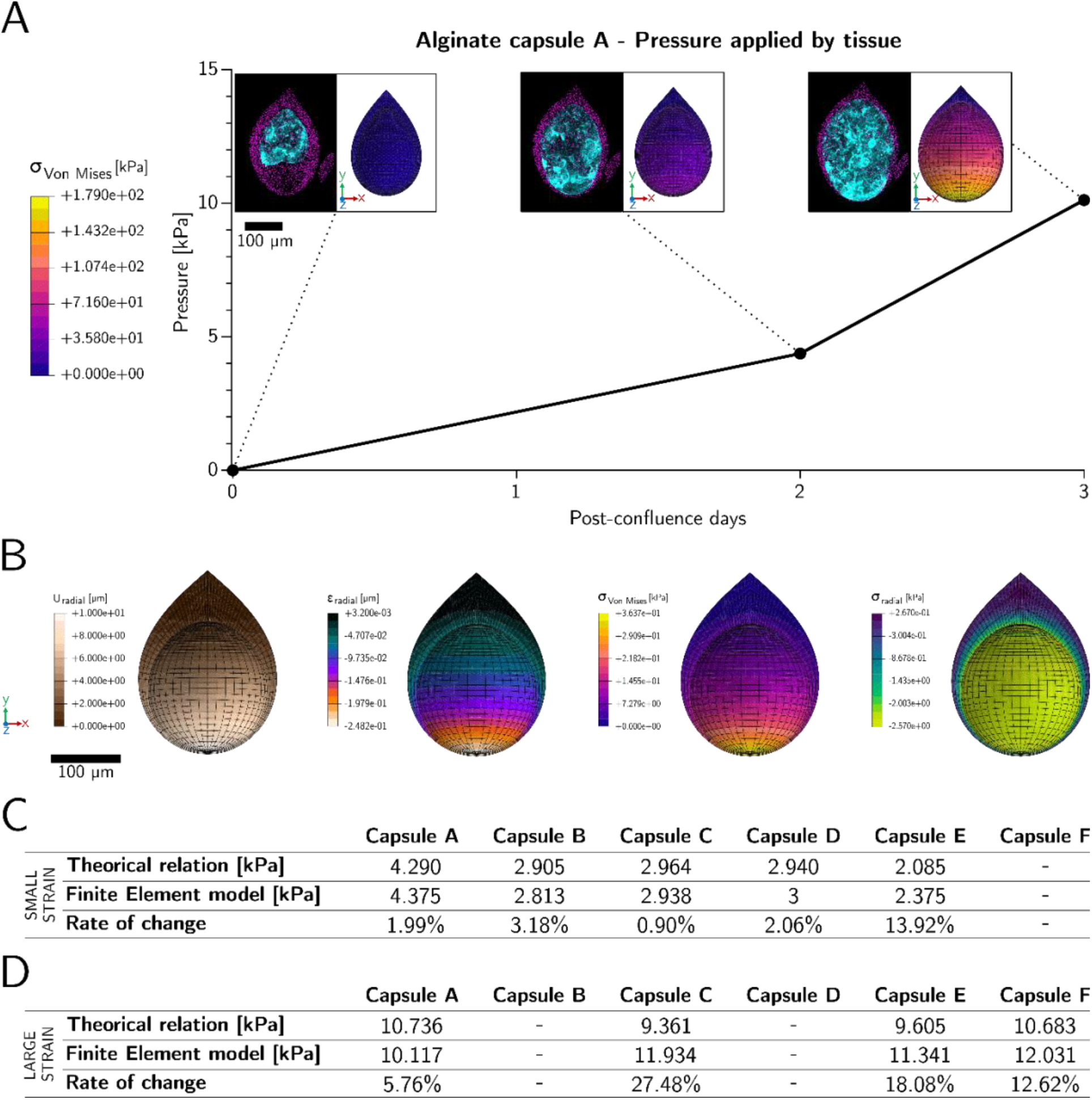
Pressure exerted by cells aggregate on the inner wall of alginate shell. (A) Pressure applied by tissue on the internal surface of the thin capsule over confluence times. The pressure unit is kPa. Day 0 is the reference day of the study, the pre-confluence times. Above the curve, there are couples of images. On the left of each couple, this is the microscopic image of the capsule A with fluorescent dots (magenta) and cellular aggregate (cyan). On the right, this is the representation of the VM stress map in the capsule, over time. (B) Types of results available using FE model: radial displacement field (µm), radial strain map, Von Mises stress map (kPa) and radial stress map (kPa). (C) Comparison table for small strain (*ε* < 10%) between pressure obtained with the theorical relation and pressure obtained with the FE model. Pressures are expressed in kPa. (D) Comparison table for large strain (10% < *ε* < 80%) between pressure obtained with the theorical relation and pressure obtained with the FE model. Pressures are expressed in kPa.

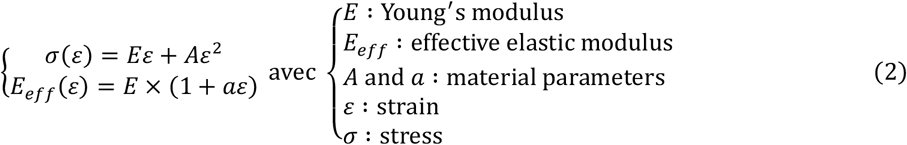

Note that *E*_*eff*_ is dependent on the strain *ε* in the regime of large deformations. The non-linearity of stress with respect to strain indicates hyperelastic behaviour; we model the material law of alginate using the Marlow law[42] (see Material and Methods section). This law is a tabulated law, calculated from pairs of values (*ε, σ*(*ε*)) obtained using the system of equations (2). The inverse method derived from these assumptions allows us to calculate, in a common reference frame, the evolution of the pressure exerted by the cells on the shell of each of the six alginate capsules. Curve 1 in Figure 3A shows this evolution for capsule A. The images above illustrate, on the left, the growth of the encapsulated tissue in confocal microscopy and, on the right, the distribution of von Mises stresses estimated by the FE model. This comparison highlights the similarity between the deformation of the FE model and that of the alginate capsule deformed by the cell aggregate. It also reveals that the stress is maximal at the bottom of the capsule, where the thickness is minimal, and zero inside the tip. As illustrated in Figure 3B, the FE model also allows for a detailed 3D analysis in the form of maps (radial displacements, radial deformation, VM stress, and radial stress) of the alginate capsule’s response to stresses. Tables 3C and 3D compare these results to the theoretical relationship, established for capsules assumed to be spherical, given by:

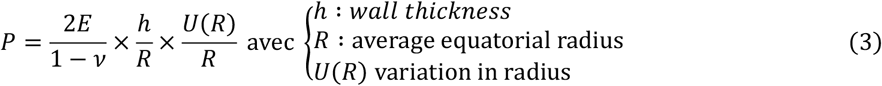

These tables distinguish between small deformations (*ε* < 10%) in Figure 3C and large deformations (10% < *ε* < 80%) in Figure 3D. To ensure the consistency of equation (3) with the values of ε in the domain of large deformations, the comparison with our model was performed by replacing the Young’s modulus *E* = 94 *kPa* with the effective modulus *E*_*eff*_. In the domain of small deformations, the numerical simulation produces results close to the theoretical formula, confirming the consistency of our model. In the domain of large deformations, the discrepancies with the theoretical values are greater, which can be explained by different geometric assumptions. Indeed, our model computes non-spherical capsules with distinct internal and external tips, leading to variability in the wall thickness. Given that our inverse method relies on 3D displacements within this wall (whose thickness is no longer constant), the obtained pressure values necessarily differ, explaining the observed discrepancies.

### 2.4. Implementation of the Young’s modulus variation with a thick alginate shell

Up to this point, our modelling was based on capsules with a thickness of less than 10 µm. We hypothesize that for thicknesses greater than 15 µm, calcium ions diffusion leads to a weaker internal cross-linking, therefore corresponding to a gradient of Young’s modulus from the outside to the inside. We are investigating whether integrating a variable Young’s modulus, dependent on the concentration of calcium ions, into our FE model would better reproduce experimental results. Initially, we demonstrate that the assumption of a constant E in the EF model is not valid with respect to experimental data. For this purpose, we will use experimental data on the position of microbeads at the pre-confluence time *T*_0_ and at the *n*_*T*_ post-confluence times *T*_*i*_. From the pre-confluence data, we select a portion of the capsule (red partition, Figure 4A1), divided into *n*_*c*_ of equal thickness and a similar number of points. The number of layers corresponds to the elements in the thickness of the FE model (e.g., 7 layers for 7 elements, Figure 4A2). This segmentation allows the separate analysis of the displacements of the different layers (internal and external) and thus provides insights into a possible variation in the Young’s modulus. The points of the inner surface are then identified and interpolated to reconstruct this surface (Figure 4A3). For each layer j, we calculate 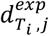, the average distance of its points to this surface at the post-confluence time *T*_*i*_ (Figures 4A4-5-6). Initially, the corresponding numerical distances, 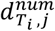, are simulated using our FE model, assuming a constant Young’s modulus of 94 *kPa* and applying the average experimental displacements of the inner surface at each time *T*_*i*_. By subtracting, for each layer j and time *T*_*i*_, the experimental and numerical distances, we obtain 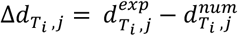. We assume that this difference is zero at the reference time 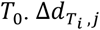 is represented in Figure 4B, where the goal is to achieve 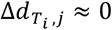 for each post-confluence curve. However, at times *T*_2_ and *T*_3_, significant deviations of -1 µm to -4 µm are observed, indicating that the FE model does not accurately describe the material law of alginate for thick-walled capsules.

**Figure 4.**
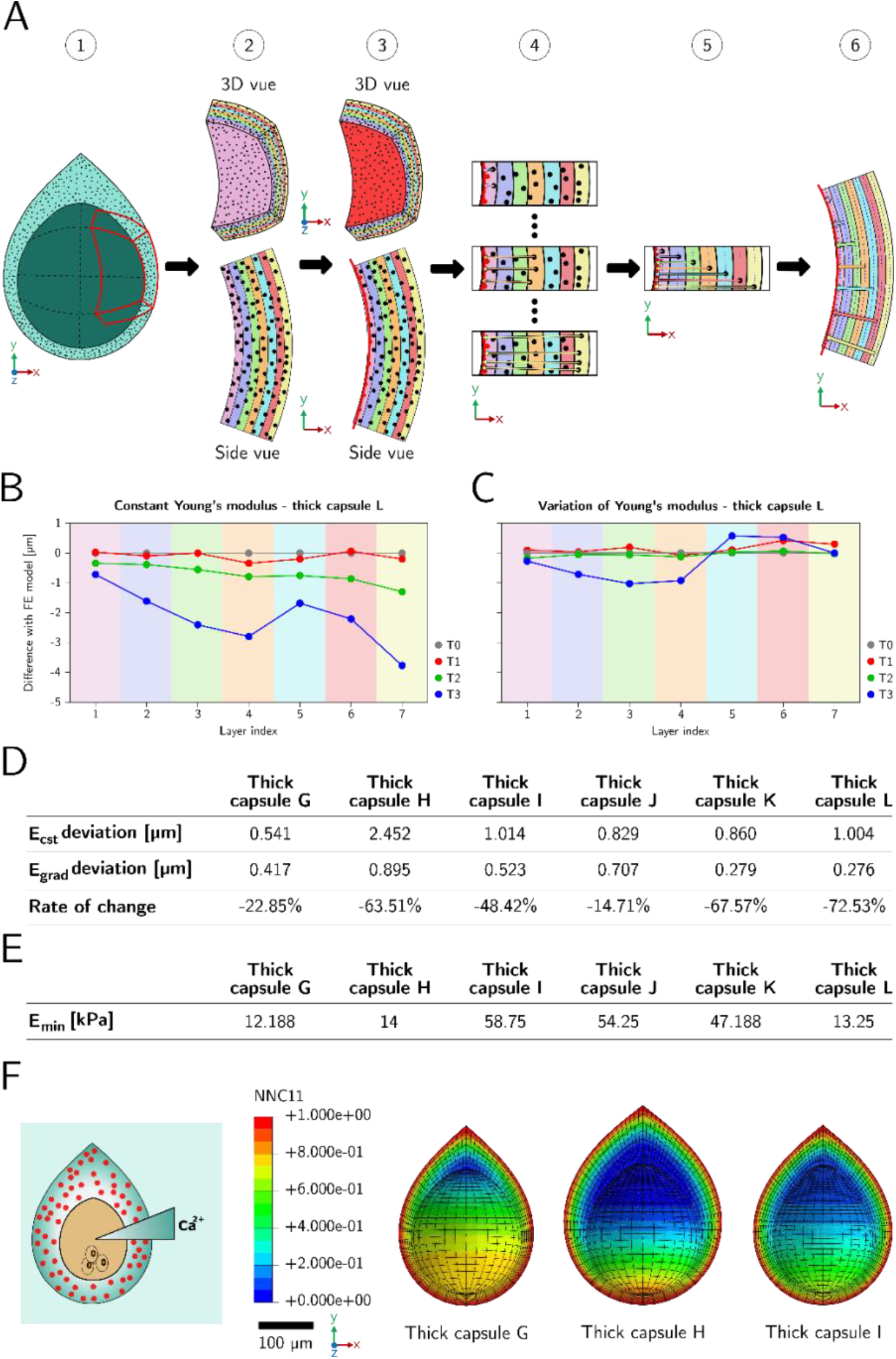
Importance of implementing a Young’s modulus variation when the alginate shell is thick. (A) Schematic of method used to obtain experimental data for the concentration gradient study. 1 – Partition selection on the experimental point cloud. 2-Layers creation on the dots partition. 3 – Inner surface interpolation. 4 – Measurement of the distance between the dots and the inner surface, layer by layer. 5 – Calculation of the average distance from each layer to the inner surface using the distances previously measured. 6 – Overview of average distances. (B) Graph showing the distance difference to the inner surface between Fe model and experimental data with constant Young’s modulus for each layer at each time. (C) Graph showing the distance difference to the inner surface between Fe model and experimental data with Young’s modulus gradient for each layer at each time. (D) Table of deviations between: constant Young’s modulus model and experimental data (first line); Young’s modulus gradient model and experimental data (second line). Third row shows the rate of decrease involved using a modulus gradient model. (E) Table of minimum internal Young’s modulus values for each thick-walled capsule. (F) On the left: schematic of calcium ion diffusion through the alginate capsule. On the right of the legend: Numerical modelling of calcium concentration profiles in different thick-walled capsules. These profiles are linked to Young’s modulus. The legend shows the ion concentration within the capsule, ranging from 0 (minimum internal modulus) to 1 (maximum external modulus).

Since the assumption of a constant Young’s modulus is no longer valid for capsules with thicknesses greater than 15 µm, we integrated a simulation of the calcium ion concentration gradient in the alginate via a mass diffusion analysis using ABAQUS (see Material and Methods). This simulation allows for a linear variation of the ionic concentration across the capsule thickness as a function of E. To adapt the diffusion profile to the considered thicknesses (>15 µm) while ensuring consistency with the modelling of thinner capsules (<10 µm) where the Young’s modulus was constant, we halt the numerical diffusion when the maximum concentration thickness reaches 10 µm. Each capsule then exhibits a distinct concentration profile, dependent on its geometry, despite identical diffusion times. Figure 4F illustrates these profiles, where the maximum concentration (in red) remains within the defined range. As we assume that *E*_*ext*_, the Young’s modulus corresponding to this concentration, is 94 *kPa*, the numerical distances 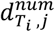, and thus the deviations 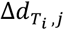, depend solely on *E*_*m*_, the Young’s modulus associated with the minimum concentration. To estimate this value for each capsule, we minimize the average deviation between the experimental and numerical distances, considering all *n*_*T*_post-confluence times and *n*_*c*_ layers:

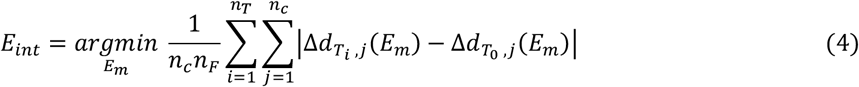

The obtained values of *E*_*int*_ are presented in Table 4E. Their heterogeneity can be explained by the differences in diffusion and geometry specific to each capsule. We then compared the FE model with a constant Young’s modulus to the one incorporating its variations by analysing the deviations for all the studied capsules (Table 4D). For the six capsules, the divergence is systematically lower with the variable modulus model, reducing the overall deviation by up to 70% compared to the constant model. In the model of Figure 4B for 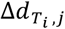, Figure 4C plots 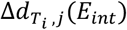, where the goal is to achieve 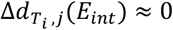 for each post-confluence curve. This graphical representation of the results shows a significantly reduced deviation between each post-confluence time and the reference time *T*_0_. Unlike Figure 4B, where the curves deviated from *T*_0_, they cluster around this reference here, indicating better agreement with the experimental data. For future studies on thick-walled capsules, we will therefore consider the Young’s modulus to be directly related to the diffusion gradient.

### 2.5. Numerical analysis to validate the addition of Young’s modulus variation in the capsule model

With the addition of the Young’s modulus variation locally modifying the material law of the alginate capsules, we replicated the analyses from Figure 2 by applying them to the FE model incorporating this variation. Similar to the sensitivity analysis presented in Tables 2B and 2C, simulations were conducted with values varying by ±30% around *E*_*ext*_ = 94 *kPa, E*_*int*_ = 50 *kPa*, and *P* = 4 *kPa*, corresponding to the extreme values in Table 5B. The results presented in Table 5B quantify the impact of each parameter on the stress. As before, the internal pressure significantly influences the stress more than the internal and external Young’s moduli. We thus draw a similar conclusion: for the remainder of the study, we will fix *E*_*ext*_ at 94 kPa and use the previously estimated values for *E*_*int*_, as their influence is negligible compared to that of the internal pressure.

The reproduction of the robustness study (Figure 2D) on the inverse method incorporating the Young’s modulus variation with respect to errors in the extraction of microbead positions was conducted with a minimum internal Young’s modulus set to *E*_*int*_ = 12.188 *kPa* (the smallest Young’s modulus value obtained previously) and a pressure of *P*^*ref*^ = 5 *kPa*. The results, presented in Figure 5C, are visualized using a violin plot illustrating the distribution of Δ*P* for each of the 100 trials performed per *σ*^2^ value. The linear regression graph allows for direct evaluation of the sensitivity to the inserted noise on the microbead positions. The slope of this regression, equal to 1e-5, represents less than 1% of *P*^*ref*^, confirming that the algorithm remains insensitive to noise in this configuration.

**Figure 5.**
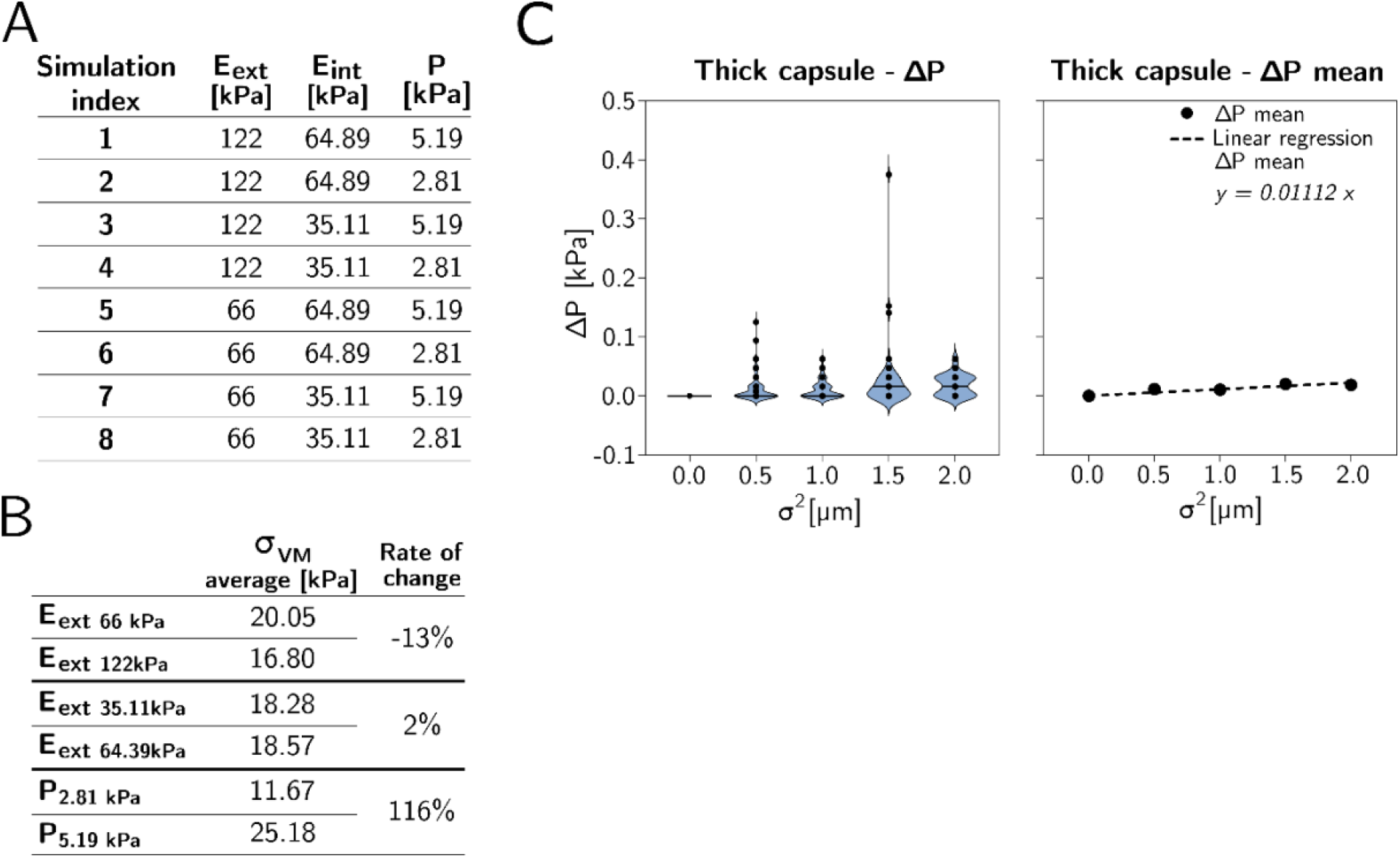
Numerical analysis on thick-walled capsule model. (A) Values of parameters used for the design experiment of mechanical parameter on results for thick-walled alginate capsule model. (C) Variation of VM stress according to Young’s modulus and pressure values for thick-walled alginate capsule model. (D) Noise sensibility of the thick-walled alginate capsule model when pressure varies. Left: violin plot of the results distribution, right: averaged values and linear regression, corresponding to the cost function sensitivity to noise.

**Figure 6.**
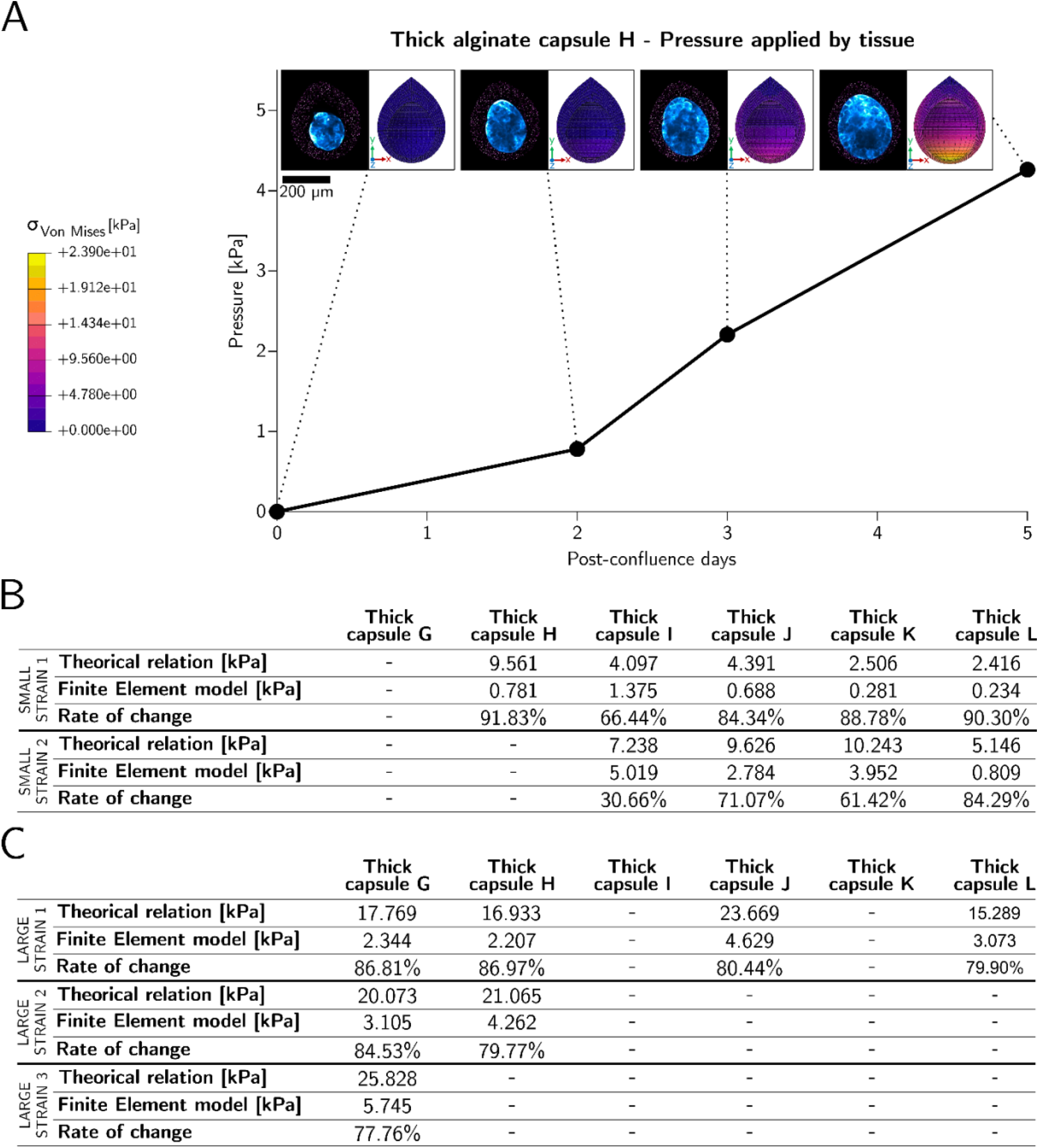
Pressure exerted by cells aggregate on the inner wall of thick-walled capsules. (A) Pressure applied by tissue on the alginate shell of the thick-walled capsule over confluence times. The pressure unit is kPa. Day 0 is the reference day of the study, the pre-confluence times. Above the curve, there are couples of images. On the left of each couple, this is the microscopic image of the capsule A with fluorescent dots (magenta) and cellular aggregate (blue). On the right, this is the representation of the VM stress map in the capsule, over time. (B) Comparison table for small strain between pressure obtained with the theorical relation and pressure obtained with the FE model. Pressures are expressed in kPa. (D) Comparison table for large strain between pressure obtained with the theorical relation and pressure obtained with the FE model. Pressures are expressed in kPa.

### 2.6. Pressure exerted on the inner wall of thick alginate shells by cell aggregates

With our model incorporating the variation of the Young’s modulus numerically validated, we extended the study conducted on thin capsules (Figure 3) to six thick-walled capsules, labelled G to L, with internal moduli corresponding to the values in Table 4E. Graph 6A shows the evolution of the pressure exerted by the cells on capsule H. As before, pairs of images positioned above the curve illustrate the growth of the encapsulated tissue at the same time points. To the right of each pair of images, the FE simulations display the Von Mises stresses induced by the cellular push. The results confirm the trends observed for the thin capsules: the stress is maximal at the bottom of the capsule and minimal at the tip. Tables 6B and 6C compare the pressure values between the FE model and the formula for thick-walled capsules[22]:

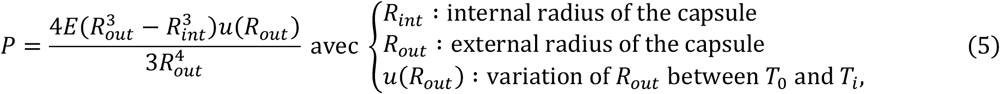

This relationship has limitations for thick-walled capsules because the assumption of constant tangential stress across the entire thickness is no longer valid. Table 6B focuses on the regime of small deformations, while Table 6C addresses large deformations. Each capsule can transition between these two regimes depending on the post-confluence time. For example, capsule G immediately enters the regime of large deformations at the first post-confluence time. Similarly, capsule H initially experiences small deformations on the first day after confluence before shifting to the regime of large deformations in the following days. In both regimes, the pressure estimated by the theoretical law differs significantly from that obtained by the FE model, with discrepancies reaching up to 90%. Specifically, the theoretical law tends to overestimate the pressure compared to the FE model. This divergence is primarily explained by the incorporation of the Young’s modulus variation in the numerical model, whereas the theoretical relation assumes a constant modulus.

### 2.7. Behavioural differences between thin and thick-walled capsules

We compare the impact of capsule thickness on the pressure exerted by the encapsulated tissue. Graph 7A shows the evolution of the average pressure for six samples of thin and thick capsules, along with their standard deviation (blue and pink areas, respectively). We calculated the average pressure values and standard deviation at each post-confluence time step. The blue area on the curve for thin capsules represents this standard deviation, while the pink area corresponds to the standard deviation for thick-walled capsules. It is immediately apparent that the tissue encapsulated in thin capsules exerts significantly higher pressure on the inner wall than the tissue encapsulated in thick-walled capsules. This provides a biological insight into these tissues, which react differently depending on the capsule thickness. This difference could be explained by the distinct elastic modulus values of the two types of capsules: cells encapsulated in thin capsules are in contact with a thinner but very rigid wall, whereas cells encapsulated in thick-walled capsules are in contact with a thicker wall, whose rigidity is low at the inner surface and gradually increases within the capsule.

The analysis reveals a significantly higher pressure in the thin capsules. This difference could be attributed to the distinct mechanical properties of the two types of capsules. In thin capsules, the cells are in direct contact with a rigid and relatively thin wall, while in thick capsules, they interact with a wall whose rigidity is lower on the inside and increases gradually towards the outside. This distribution of the Young’s modulus could influence how cells exert pressure on the shell: they may exert less pressure on thick-walled capsules, causing less deformation to the alginate. Figure 7B illustrates these differences in cellular behaviour: the Von Mises stress is significantly higher in thin capsules than in thick capsules for the same post-confluence time. We previously showed that thick capsules exhibit a variation in Young’s modulus across their thickness, whereas thin capsules can be assumed to have a constant modulus. We then wanted to compute the thin capsules by applying a similar gradient, which logically would lead to an almost constant Young’s Modulus value considering the very small distance for the calcium ions to diffuse through. Figure 7C highlights these differences by illustrating the distribution of gradients for an identical diffusion time. The analysis reveals that this gradient is much less pronounced in thin capsules, which can be attributed to their smaller thickness, facilitating more homogeneous diffusion compared to thick capsules. For thick capsules, this gradient shows that the maximum diffusion zone (in red) extends approximately 10 µm, comparable to the thickness of the bottom of thin capsules. If the maximum rigidity is associated with this 10 µm zone in thick capsules, then the absence of a diffusion gradient in thin capsules appears consistent with the observations in the figure.

**Figure 7.**
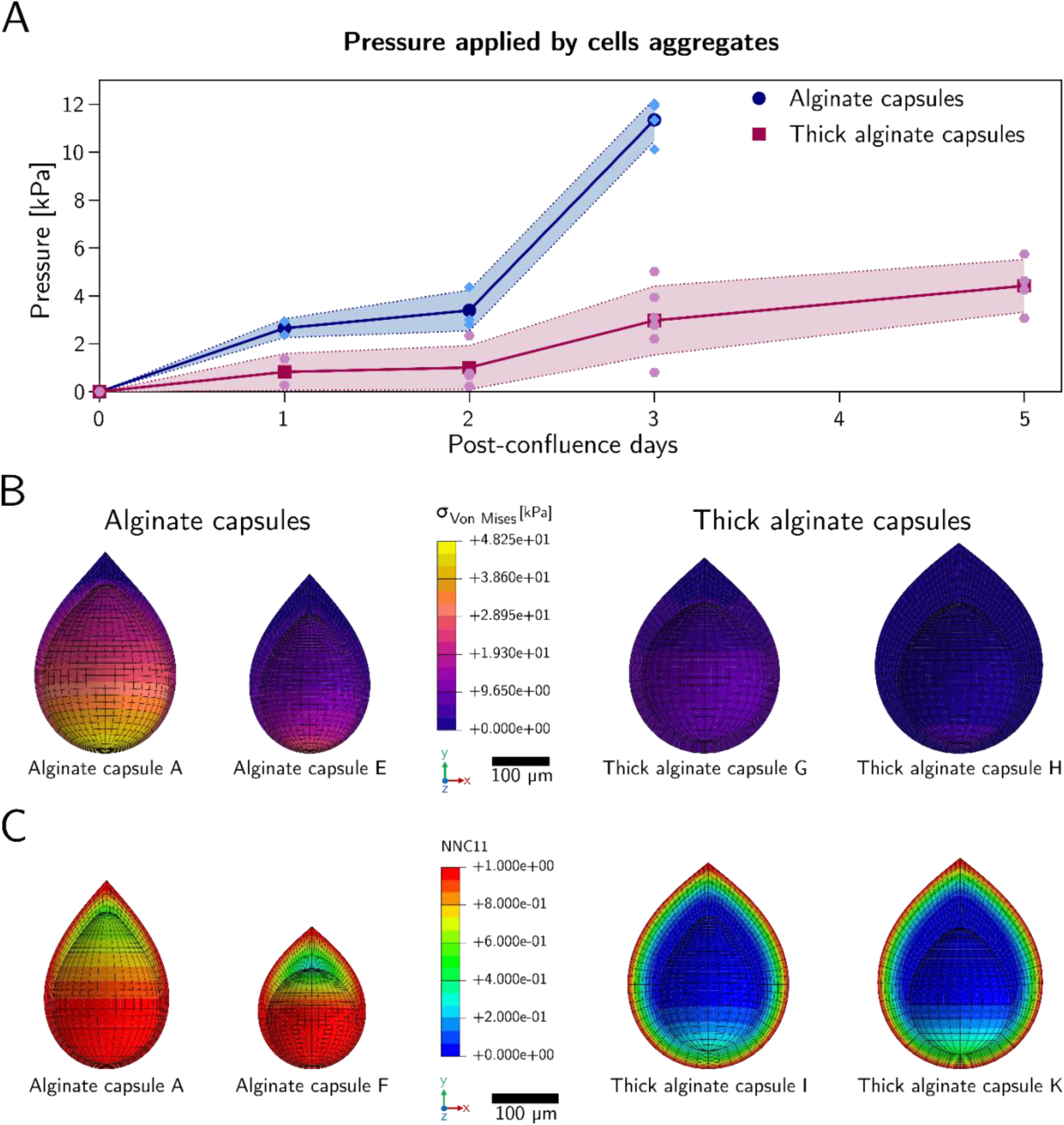
Differences between thin capsules and thick capsules. (A) Averaged pressure evolution over post-confluence times for thin and thick-walled capsules. The blue surface represents the standard deviation of thin capsules (6 capsules) pressures. The rose surface represents the standard deviation of thick-walled capsules (6 capsules) pressures. (B) Differences in Von Mises stress between thin capsules (left) and thick-walled capsules (right). (C) Differences in calcium ion concentration gradient between thin capsules (left) and thick capsules (right).

## 3. Materials and methods

### 3.1. Caco-2 cells culture and alginate encapsulation

Caco-2 (DSMZ, acc169, gift from Audrey Ferrand) cells were cultured at 4000 to 8000 cells/cm2, passaged every 3 to 4 days, in DMEM (High Glucose, with L-Glutamine, with Sodium pyruvate, L0104-500, Dominique Dutscher) with 10 % FBS (FBS-16A, Capricorn Scientific), 1 % Penicillin-Streptomycin (P06-07100, Pan Biotech), and 1 % MEM non-essential amino acids (11140-050, Gibco), at 37°C and 5% CO2.

As previously described[22,23], we used the Cellular Capsule Technology (CCT) to incorporate Caco-2 cells in a basement membrane extract (BME) matrix (Cultrex BME001, R&D Systems) surrounded by a 3D spherical shell of alginate (AGI, I3G80). Unless stated otherwise, we use a cell formulation of 100 000 Caco-2 cells in 200µL of DMEM with 50% [v/v] BME maintained at 4°C during encapsulation. Briefly, a microfluidic chip coextrudes the cell formulation inside an outer jet of alginate 2% [w/v] supplemented with 1% red-fluorescent microbeads (Sicastar®-redF plain 1µm), both separated by solution of sorbitol (300mM, 85529, Sigma). The stream breaks into droplets that are collected in a CaCl_2_ bath (100mM, 7774-34-7, Themo scientific) gelling the alginate into a wrapping capsule. The cell-containing capsules are then cultured in suspension in cell culture medium at 37°C and 5% CO2.

### 3.2. Experimental determination of alginate Young’s modulus in capsules

To experimentally determine the Young’s modulus of capsules in the case of a thin thickness, we conduct an osmotic swelling study. For this purpose, we consider an alginate capsule in which a Dextran solution is encapsulated, immersed in a Dextran bath. At the beginning of the experiment, *T*_0_, the internal and external Dextran concentrations of the capsule are identical (*c*_0_) and the average radius of the capsule is denoted as *R*_0_. The external concentration is decreased by dilution, leading to a concentration decrease at *T*_1_ (decrease from *c*_0_to *c*). Since the capsule is permeable to water but not to Dextran, water diffuses into the capsule, causing it to swell until the osmotic pressure Π_0_ is balanced, which increases the initial radius by Δ*R* = *R* − *R*_0_ and decreases the initial thickness *h*_0_. It is then possible to calculate the Young’s modulus using the following relation[22]:

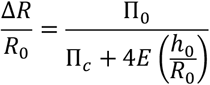

With 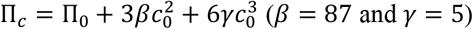. We perform this protocol on 8 different capsules and obtain an average Young’s modulus of *E* = 94 *kPa* ± 28 *kPa*. This value will be used as the constant Young’s modulus for thin capsules and as the external Young’s modulus for thick capsules

### 3.3. Virtual Model geometry

External and internal curves of the alginate capsule are based on the Booth Lemniscate’s 2D equations:

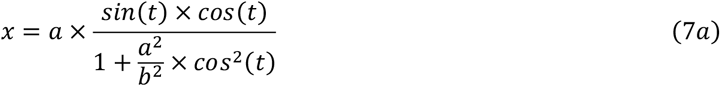

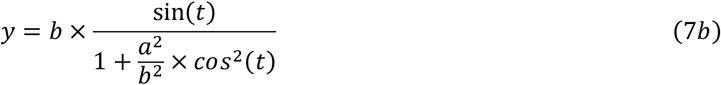

Where *a* is a parameter depending on the half-width, at the widest place of the lemniscate, *b* the half-height of a lemniscate defined on *[0,2π]* and *t* the angle parameter defined on *[0,π]*, in order to obtain a quarter of lemniscate.

A revolution operation is applied along the y axis to obtain the capsule. Tip of internal lemniscate is smoothed to avoid geometrical irregularities. The dimensions of the capsule (internal height, internal half width, external height and external half width) are obtained from the experimental point cloud. External dimensions are directly taken from the dots positions. Internal dimensions are obtained by selecting the dots belonging to the internal surface, using an alpha-shape algorithm[43]. The thickness at the capsule bottom is also obtained directly on the capsule point cloud.

### 3.4. Finite Elements choice

The capsule finite elements model is composed of 20-node hexahedrons (C3D20H) and 15-node prisms (C3D15H – wedge elements). Hexahedron elements are the best-performing volumetric elements. Prism are used to make connections and facilitate meshing where hexahedron cannot be used (top and bottom of the capsule). Quadratic elements are used because this gives a better representation of the capsule curved shape. Moreover, this enables a better analysis of the stresses distribution. A hybrid formulation is chosen because alginate is considered as an incompressible material (the volume cannot change with loading). This avoids potential convergence problems during the simulation. We conducted a mesh convergence study to determine the mesh offering the best compromise between results accuracy and simulation cost. This analysis is based on various criteria: Von mises maximal Stress, *σ*_11_max, …, and the Aspect Ratio Value. The aim is to find a compromise between results accuracy on each stress, calculation time and satisfactory elements aspect ratio.

### 3.5. Choice of the material law

Alginate is a biocompatible, incompressible and elastic material. To study its material law, we use the following data[22]: Poisson’s ratio is equal to 0.5 and elasticity depends on the deformation rate: for deformation below 10%, alginate has purely linear elastic behaviour and above 80% deformation, plasticity appears. For deformations between these two thresholds, alginate presents a non-linear stress-strain response and thus a non-linear elastic behaviour (eq. 2).

We use a hyperelastic law to represent numerically the material behaviour. To use the software Simulia Abaqus, some information is needed to extend the unidirectional law (eq.2) in 3D. We use the Marlow hyperelastic law [42], which considers that strain energy potential is independent of the second deviatoric invariant. The deviatoric behaviour is defined by *W*_*dev*_, which is characterized by test data like uniaxial traction test[44]. The volumetric behaviour is defined by *W*_*vol*_ if the material studied is compressible. The strain energy potential[45] is thus defined by:

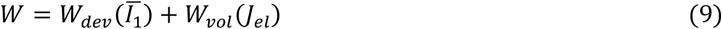

Where 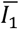 is the first deviatoric invariant and *J*_*el*_ is the elastic volume ratio :

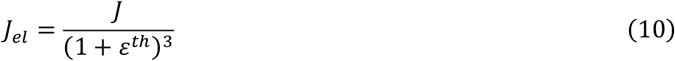

Where *ε*^*th*^ is the linear thermal expansion strain and *J* is the total volume ratio. In our case, alginate is an incompressible material with a Poisson’s ration equal to 0.5 so there is no volume variation during analysis. The strain energy potential is therefore defined solely by *W*_*dev*_ :

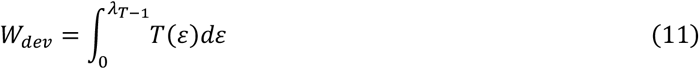

Where *T*(*ε*) is the nominal uniaxial stress obtained by single uniaxial tension test and *λ*_*T*_ the uniaxial stretch. The data table is constructed from equation (2a). The minimum value of deformation rate is 0.1, because when deformation rate is lower, alginate behaves like a purely elastic material. The maximum value of *ε* is 0.8, above plasticity and fracture appear. The value taken for the Young’s modulus, *E*, is 94 *kPa* ± 28 *kPa*, which is determined with the osmotic pressure measurement with Dextran on experiments.

### 3.6. Boundary conditions and loads

Numerical boundary conditions are not the same than experimental ones. Experimental capsules rest on the bottom of a calcium bath with flat support linkage. However, this flat support is achieved randomly, with the capsule tip pointing upwards or downwards. At post-confluence times, when capsule sweels due to pressure exerted by encapsulated cells within it, boundary conditions may change slightly. Swelling can cause a rotation of the capsule and changing the initial flat support into a different flat support. However, these rigid body movements are corrected during the experimental dots tracking with *Fiji – ImageJ* software. Thus, this has no impact on the final displacements. Numerical boundary conditions need to be set up to take possible experimental configurations in account while maintaining the same boundary conditions over time to respect the modelling assumptions. To achieve this, the capsule is fully modelled, and the applied boundary conditions allow isostatic embedding. For this, the X-axis translation X axis is blocked, as is Z-axis. The Y-axis translation is blocked on the capsule tip. With these boundary conditions, the capsule can be freely deformed by the internal pressure, without being constrained by these conditions. However, they are very different than those implemented during experiments. To overcome this problem, numerical data are registered using our dedicated python algorithm. Rigid transformations are applied. This has no impact on deformation or on dots positions, since they are only translated or oriented without changing scale. About experimental load, there is no local dots displacement until cells have filled the capsule internal cavity. So, pressure can be modelled by a uniform internal pressure applied at confluence time.

### 3.7. Mass diffusion analysis

In order to represent the calcium ion diffusion through alginate capsule, a Mass diffusion analysis, on Abaqus, was set up. This analysis models the transient diffusion of one material through another one. To use it, it is necessary to use mass diffusion finite elements, so C3D15H become DC3D15 and C3D20H become DC3D20. The number of elements and nodes remains the same as for mechanical analysis. The diffusion problem is define based on the requirement of mass concentration for the diffusing phase:

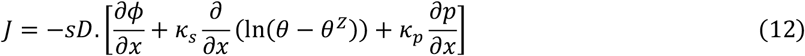

where *D* is the diffusivity; *s* the solubility; *ϕ* the normalized concentration; *κ*_*s*_ is the “Soret effect” factor; *θ* is the temperature; *θ*^*Z*^ is the absolute zero value on the temperature scale being used and *κ*_*p*_ is the pressure stress factor. In our case, temperature and pressure stress factor are not take into account so the relation become:

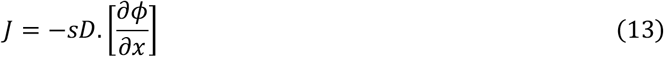

As alginate is 99% water, characteristics used to simulate calcium ion diffusion through alginate are those of water (Diffusivity = 2.895 × 10^−9^*m*^2^*s*^−1^ at 35°C)[46]. Calcium concentration in alginate at initial step is equal to zero, except on the external surface of the capsule, where the concentration is equal to one because capsule is immersed in calcium bath. A surface concentration flux is apply on the outer surface, inwards capsule. Mass diffusion analysis gives as result the evolution of calcium concentration in the alginate capsule. This concentration is between 0 (minimal concentration) and 1 (maximal concentration). To obtain a physical concentration profile, it is necessary to choose an increment to be retained for future analysis. The chosen increment should have a profile where the calcium concentration is highest on the outer surface of the capsule over a thickness of 5 to 10 µm. This ensures consistency with previous osmotic pressure tests. The chosen increment is the same for all thick capsules because time spend in the calcium bath is the same for all capsule. Calcium diffusion time is the same for all, but each have a different concentration profile, due to thickness, height and width differences between each capsules. It is then possible to consider this calcium concentration profile as a gradient of mechanical property. Where concentration is highest, the mechanical property is highest and vice versa. As the calcium ions diffusion makes alginate capsules more rigid, it is decided to treat this concentration gradient as a young’s modulus gradient.

## Conclusion and outlook

In this work, we presented a FE model of the extracellular environment of a tissue, based on experimental data integrated into an inverse method. This method proved to be robust and reproductible, as demonstrated by numerical studies conducted on different types of models. Our FE model coupled with the inverse method allows to determine the pressure exerted by the cell aggregate on the capsule and the pressure exerted by the capsule on the cells, thanks to the principle of efforts reciprocity. Additionally, we can obtain the spatial distributions and intensity of stresses within the capsule in three dimensions. The quality of the FEM of thin capsules is confirmed by the small discrepancies between the pressure results obtained with the theorical formula for small deformations and those obtained numerically. The larger discrepancies observed for large deformations are explained by the consideration of the non-linearity of the alginate material in the numerical model. In the case of thin capsules, the material is considered isotropic and homogeneous due to the complete cross-linking of the alginate capsule in the calcium bath. After validating the model under these standard conditions, we studied the case where the capsule wall is thick. We then hypothesized the incomplete cross-linking of the alginate in the innermost side, which would induce a gradient of mechanical properties. We numerically verified this hypothesis using experimental data related to thick-walled capsules, through minimization of the discrepancy between the numerical model and results from the biological model. After defining these material laws for the different types of capsules, we were able to use the implemented inverse method to determine the intensity of the pressure applied by the cells to the inner surface of each capsule. Deviations from the existing theoretical relations for thick-walled capsules[22] are significant, which is consistent since these relations only consider the case where the Young’s modulus is constant throughout the capsule. However, taking this gradient into account is not always necessary, as demonstrated by its application to thin-walled capsules with or without a gradient of mechanical properties are very close and in the same order of magnitude. Regarding the calculated pressure, we noticed a difference in pressure values between tissues encapsulated in thin-walled capsules and thick-walled capsules, which induces a difference in behaviour. Indeed, for thin capsules cells exert significant pressure due to the high rigidity of capsules. For thick-walled capsules, we observe a different behaviour, as the pressure exerted by cells on the capsule and reciprocally, is gradual. So far, we have not considered irregularities such as hernias or variations in thickness along the revolution axis used for model construction. The FE capsule model being a first step in obtaining the mechanical parameters for a multi-material micro-tissue. We model here only the surrounding environment, but it would also be insightful to embark a cellular numerical model[36] inside the capsule model provided in this study. This framework would not only be applicable to model tumour confinement and mechanics but also to model the development of embryos. As it happens, in many species, the embryo firstly develops protected by a transparent shell before hatching, the zona pellucida, chorion or vitelline membrane for mammals, zebrafishes or xenopus respectively[47–49]. [50,51] Stressing the need of providing numerical twins in biology. Often, these are based on a different model from FEM. They use one[50] or more particles[51], governed by simplified yet biologically relevant mechanical laws, to represent a cell. Although these approaches require high computational time and are not well suited for modelling continuous materials such as alginate, FEM provides a mechanically rigorous framework while maintaining reasonable computation times.

## Acknowledgements

We wish to thank Julien Laussu, Deborah Michel, Léa Magne, Dimitri Hamel, Pierre Nassoy and Adeline Boyreau for critical discussions and suggestions on this work. We also thank BIC and Plateforme Voxel for technical assistance.

## Notes

### Competing Interest Statement

The authors have declared no competing interest.

